# A Tailored Approach To Study Legionella Infection Using Lattice Light Sheet Microscope (LLSM)

**DOI:** 10.1101/2022.03.20.485032

**Authors:** Xiyu Yi, Haichao Miao, Jacky Kai-Yin Lo, Maher M. Elsheikh, Tek Hyung Lee, Chenfanfu Jiang, Yuliang Zhang, Brent W. Segelke, K. Wesley Overton, Peer-Timo Bremer, Ted A. Laurence

## Abstract

*Legionella* is a genus of ubiquitous environmental pathogens found in freshwater systems, moist soil, and composted materials. More than four decades of *Legionella* research has provided important insights into *Legionella* pathogenesis [1]. Although standard commercial microscopes have led to significant advances in understanding *Legionella* pathogenesis [2,3], great potential exists in the deployment of more advanced imaging techniques to provide additional insights. The Lattice Light Sheet Microscope (LLSM) is a recently developed microscope for 4D live cell imaging with high resolution and minimum photo-damage [4]. We built a LLSM with an improved version for the optical layout with two path-stretching mirror sets and a novel Reconfigurable Galvanometer Scanner (*RGS*) module to improve the reproducibility and reliability of the alignment and maintenance of the LLSM. We commissioned this LLSM to study *Legionella pneumophila* infection with a tailored workflow designed over instrumentation, experiments, and data processing methods. Our results indicate that *Legionella pneumophila* infection is correlated with a series of morphological signatures such as smoothness, migration pattern and polarity both statistically and dynamically. Our work demonstrates the benefits of using LLSM for studying long-term questions in bacterial infection. Our free-for-use modifications and workflow designs on the use of LLSM system contributes to the adoption and promotion of the state-of-the-art LLSM technology for both academic and commercial applications.

## 1. Introduction

*Legionella* is an omnipresent bacterial genus found in both natural environment such as lakes, moist soil and composted materials, and man-made environment such as the urban freshwater pipe systems [1,4–7]. Within the *Legionella* genus, the *Lg. pneumophila* strain is known to cause *Legionnaire’s disease*, a rapid pneumonia with 0.1-14% attack rate [8] and up to 80% case-fatality rate within 2 to 8 days post exposure [9]. After infection, the *Lg. pneumophila* bacteria translocate more than 300 effector proteins into the host cell through a Type-IV secretion system (T4SS) [10], evade the host cell defense mechanism by hijacking the physiology of the host, and create the *Legionella Containing Vacuole* (LCV) as a safe niche for its own intracellular survival and replication [1]. The presence of *Legionella* bacteria in both natural and urban environments, and its potential to cause the fatal *Legionnaire’s disease*, pose a long-standing threat to the public health.

More than four decades of study on *Legionella* pathogenesis has delivered important insights about its infection, intracellular survival and replication, as well as its alteration of the signaling pathways involved in the responses of the infected cells. A recent review article has summarized 50 *Legionella* effector proteins investigated in molecular detail [1]. Over 1700 proteins were identified to be relevant to the *Legionella* infection based on genomics and proteomics studies [11,12], and the transposon insertion sequencing technique has been adopted to study the long-range interactions of genes as well as the gene expression profile involved with *Legionella* infection [13]. While the previous studies were generally inspired by the comprehensive profiling of the relevant genes, proteins, and their transcription profiles, a few recent studies were inspired by the phenomenological signatures uniquely observed through imaging techniques. In particular, bright field and fluorescence microscopy were used to study the mobility of the host cells influenced by a *Legionella* effector protein (LegG1) [14]. Fluorescence microscopy also contributed to the discovery of the preferential localization of LCV at the exit end of the ER [15]. Additionally, using fluorescence microscopy for live cell imaging, scientists have observed fragmented mitochondria inside the host cells after *Legionella* infection [16] and further discovered that *Legionella* effector proteins hijack the membrane fission and fusion processes on the host cell mitochondria [3]. Furthermore, time lapse confocal microscopy was used to observe heterogeneity of the *Legionella* phenotypes in biofilms and infected cells and unveiled the function of the *Legionella* quorum sensing system and its essential role in the formation of a subpopulation of virulent *Legionella* persisters [17]. In summary, the utilization of commercial fluorescence microscopes has successfully complemented the biomolecular approaches by providing direct observations of labeled cellular structures with spatial-temporal information. Such success suggests great potential in the deployment of more advanced imaging techniques to provide extra insights through direct observations that are previously inaccessible.

The lattice light sheet microscope (LLSM) [18] is one of the recently invented fluorescence microscopes that capitalizes on cutting-edge techniques, such as light sheet microscopy and super-resolution fluorescence microscopy, and offers unprecedented fast 4D imaging of live samples with minimum photodamage. It is a compelling tool to expand the range of explorations for the studies of bacterial infection. To date, LLSM has not been used for the study of bacterial infection to the best of our knowledge [19]. In our experience of applying LLSM for bacterial infection study, we found three major challenges. The first is to maintain the stability, robustness, and the imaging throughput of the microscope to support day-to-day imaging needs. Such challenge limits the microscope accessible to a small community requiring optics expertise for daily maintenance. The second is the design of experiments to access the unique capabilities of the microscope, and at the same time deliver biological insights about bacterial infection that are unattainable with other imaging modalities. The third challenge is the intensive postprocessing of large amounts of data with adequate pace to support the experimental explorations and the iterations required in experimental designs.

To overcome these three challenges, it is essential to bridge the knowledge and expertise between microscopists, biologists, and data scientists, and create a tailored workflow for each specific application scenario, which requires long term and deeply integrated collaborative effort across the three domains of expertise. This could be challenging for small or new research teams due to resource limitations, and poses barriers for collaborations with imaging centers due to the challenges involved with the required long-term close collaborations. The tailoring process requires innovative technical development. We believe sharing the tailored designs will accelerate the effective adoption of LLSM for the relevant application domain, and is beneficial for the both the academic research teams and the commercial providers and users of LLSM.

In this work, we developed a series of innovative approaches optimized over instrument operation and maintenance, biological experiment design, and data processing, and created a tailored workflow to use lattice light sheet microscope to study *Legionella pneumophila* infection, which can be adapted for other research topics with additional inputs. First, based on the original design licensed from HHMI, we built the LLSM with an improved version of optical layout and a novel design for aligning the galvanometer scanners, and achieved months of stable imaging operations without requiring extensive manual optical adjustment (section 2). Next, we developed a workflow to study the phenotypic effects of *Legionella* infection on macrophage cells using the RAW264.7 mouse macrophage cell line as a model. The data acquisition schemes are designed for the unique niche of observations that LLSM can offer and at the same time provide biological insights relevant to the topic of interest (section 3). Additionally, we developed a data processing workflow harnessing high performance computing (HPC) to efficiently process the data and generate reports to support the fast iterations required in the study. Our data processing scripts are also adaptable for personal computers when HPC isn’t available (section 4). With our workflow, we achieved the throughput of imaging 267 cells over 20 imaging sessions on 25 samples with 13.1% outlier rate; each dataset is a movie of one cell imaged for approximately 26 minutes. Lastly, we characterized the correlation of infection with the morphology and migration of the infected cells (section 5). Our work deploys LLSM in the study of bacterial infection and yields results to motivate the follow-up detailed investigations into the relevant biomolecular mechanisms. Our approaches are designed to be attainable for small research groups and is transferrable for well-equipped and staffed imaging centers to facilitate effective collaborations. We expect our work to be transferrable to the general domain of infectious disease study.

## 2. Instrument

### 2.1. Optical layout

We expanded the optical path from the original LLSM design onto a 3’-by-5’ optical table. As shown in Figure 1, the path contains a group constructed on the optical table (horizontal plane), and another group constructed on a 0.5-inch-thick breadboard vertically mounted on the optical table (vertical plane). We kept the specifications of the optical elements the same with the original design [18], and reconfigured the beam path to have mostly 90° reflective angles. The excitation objective is custom made by SPECIAL OPTICS (Model 54-10-7@488-910nm) and has NA of 0.66, and detection objective is purchased from Nikon (CFI75 Apo 25XC W) and has NA of 1.1. To facilitate the alignment and maintenance, we added two beam-path-stretching modules (PS1 and PS2) inspired by the eSPIM design [20] to allow for independent adjustments on the length of the beam path that is critical for aligning the lens relay system (L1 to L7 shown in Figure 1). We also designed a *Reconfigurable Galvanometer Scanners* (*RGS*) module (annotated in Figure 1 and shown in Figure 2(a)) to facilitate the precise positioning of the galvanometer scanners into the conjugated back focal planes (CBFPs), which will be explained below in detail.

**Fig. 1.**
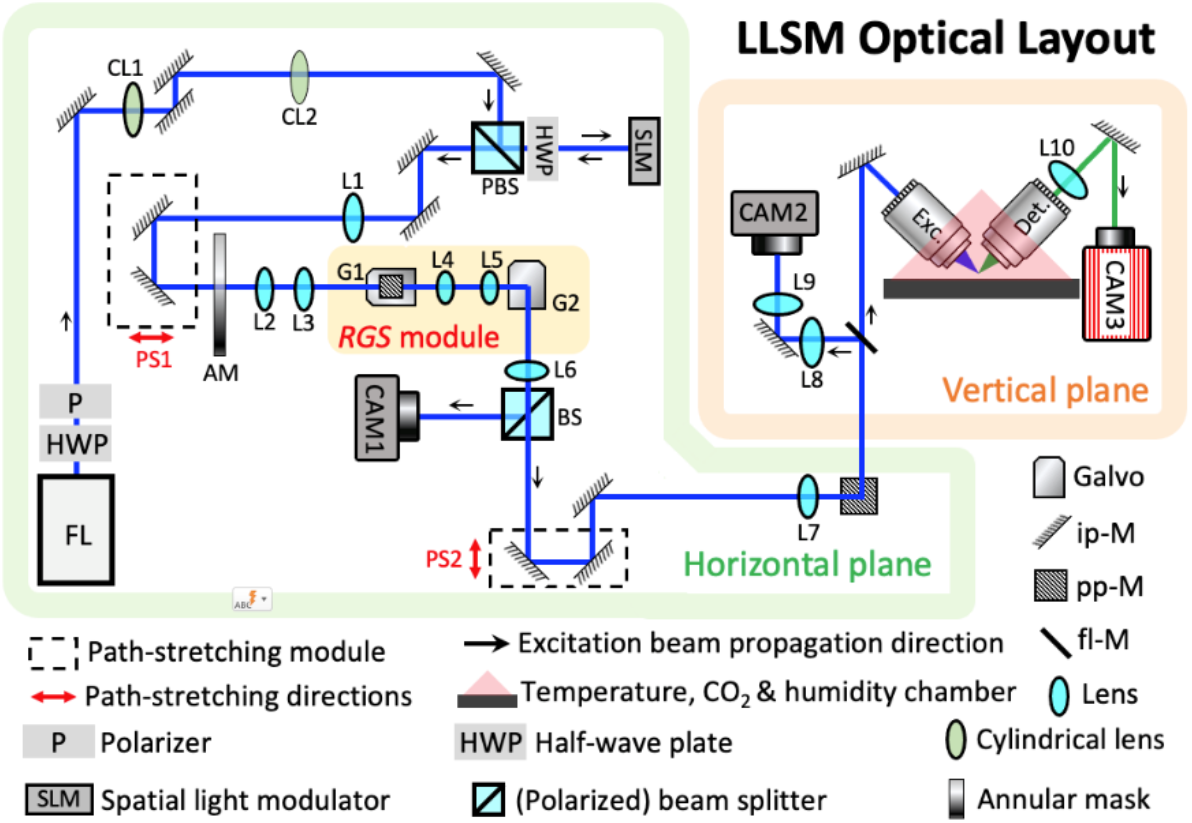
Modified optical layout of LLSM. We expanded most of the optics specifying the excitation path of LLSM onto a 3’×5’ optical table (horizontal plane) as compared to the original design while keeping the optical elements the same. The sample handling area are maintained the same on a breadboard vertically mounted on the optical table (vertical plane). FL indicates a fiber launch system to couple the single mode fiber output from a combined laser box (Oxiuss) into free-space optics. 6 excitation beams are available in our system with wavelengths of 405 nm, 488 nm, 515 nm, 561 nm, 593 nm, and 647 nm. “Galvo” indicates galvanometer mirror. “ip-M” indicates mirrors that deflect the laser beam in-plane marked as either horizontal or vertical, similarly “pp-M” indicates the mirror that deflect the laser beam in the direction perpendicular to the plane. “fl-M” indicate a flippable mirror, and CAM1 and CAM2 are inspection cameras conjugated to the imaging plane and back focal plane of the excitation objective respectively. CAM3 is the camera for imaging data acquisition.

To offer precise translational scanning of the light sheet in the focal plane of the excitation objective, the galvanometer scanners need to be placed in the CBFPs created by the 4f system with short focal length (25 mm) lenses, the space restrictions and the multiple coupled degrees of freedom pose challenges for the alignment and adjustment. The core concept of *RGS* is to offer precisely reproducible configurations of the galvanometer scanners (G1 and G2) between (a) a *normal configuration* and (b) an *alignment configuration* where the center or the edge of the galvanometer mirror is placed at the center of the optical axis respectively. The key is to set G1 and G2 into the *alignment configuration* and bring the mirror edge into sharp focus in the inspection camera (CAM2) that is imaging the CBFP.

Figure 2(a) provides an illustration of the *RGS* module. We mount each galvanometer mirror in a rotational cage mount (CRM1LT, Thorlabs Inc.) through a custom-made adaptor. The cage mount is further constructed into a 30 mm cage system with extra rods and cage plates (Thorlabs Inc.) and mounted on a XYZ-stage to facilitate precisely reproducible translational adjustments. When constructing the LLSM, we first set the *RGS* in the *normal configuration* where the beam is centered at the galvanometer mirrors, and position all the reflective optics to align the laser beam through the optical axis of the excitation objective, up to this point, the laser beam defines the optical axis for all the lenses. We then align the lenses along the optical path starting from the excitation objective following common optical alignment practices. The novelty in our design involves steps relevant to the *RGS* module and the path-stretching modules (PS1 and PS2) that starts from the positioning of L6, which will be discussed in detail below.

**Fig. 2.**
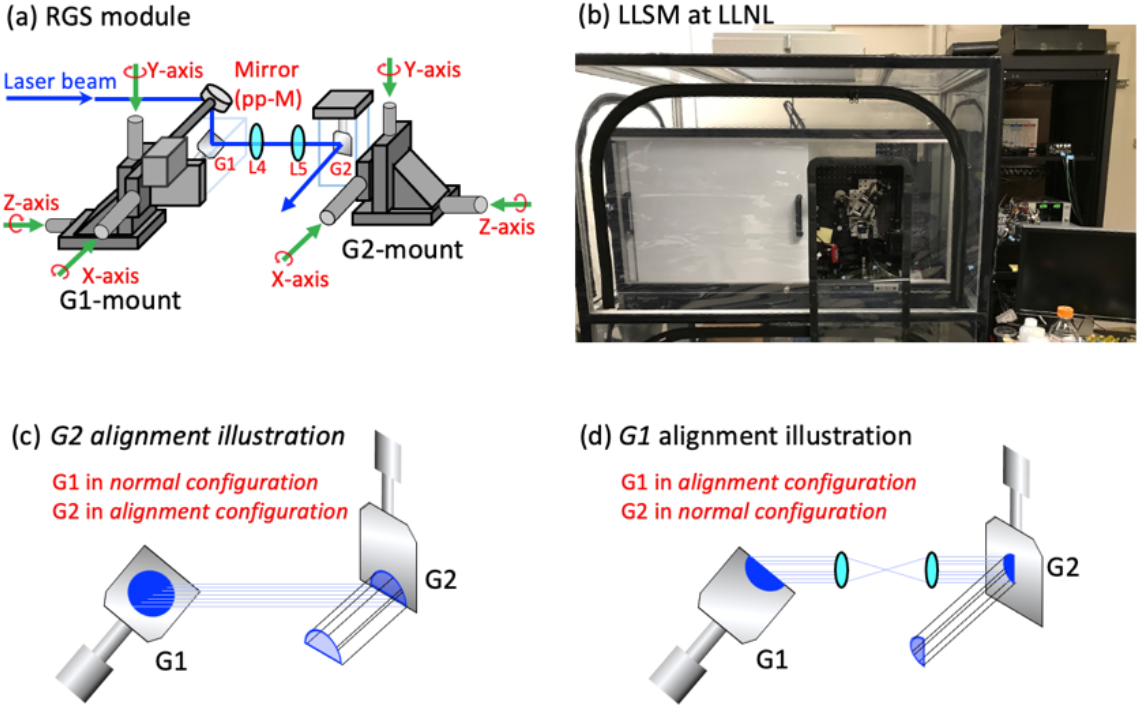
RGS module and the LLSM at LLNL. (a) shows the *Reconfigurable Galvanometer Scanners* (*RGS*) module. The *RGS* contains two galvanometer mount (G1-mount and G2 mount) each has a galvanometer (G1 and G2 respective) mounted on a XYZ translational stage through the 30 mm cage system (Thorlabs Inc.). L4 and L5 indicates the two lenses with 25 mm focal length in LLSM, and G1-module contains a mirror (pp-M) that directs the beam perpendicular to the resident plane of this module. (b) shows a photograph of the Lattice Light Sheet Microscope at Lawrence Livermore National Laboratory. In the picture, we can see the LLSM setup is covered by two layers of enclosures. The inner layer is the laser enclosure that covers the entire optical table. The outer layer is a transparent enclosure, which is the bioBUBBLE system that provides BLS-2 containment. (c) shows the *RGS* configuration for G2 alignment, where the beam is cropped by G2 and the edge of G2 would be in focus in CAM2. (d) shows the RGS configuration for G1 alignment, where the beam is cropped by G1 and the edge of G1 would be in focus in CAM2. Note that the positioning of G2 is performed before the positioning of L4 and L5, while G1 is adjusted after L4 and L5 are in position. The demonstration of CAM2 inspections while switching a galvanometer mirror between the “alignment configuration” and “normal configuration” is shown in visualization 5.

### 2.2. RGS alignment

As shown in Figure 1 and Figure 2(a), when positioning L6, we have to ensure that L6 and L7 together form an asymmetric 4f system, and at the same time the CBFP near L6 is aligned to the position of G2. However, two degrees of freedom are coupled in the adjustment. First, changing the axial position of L6 changes the length of the beam path between L6 and L7 that needs to be compensated. Second, changing the axial position of G2 would shift the beam path off from the pre-aligned directions, which could require repositioning of multiple reflective optical elements. To decouple the degrees of freedom, we implemented the path stretching module (PS2) that contains a group of mirrors mounted on a 1-axis translational stage adjusted by a manual actuator (Newport) to allow for independent adjustment of path length between L6 and L7 without moving other elements in the optical layout. As shown in Figure 2(a) and Figure 2(c), we fix the axial position of G2, set G2 into the *alignment configuration* by adjusting the Y-axis in the G2-mount shown in Figure 2(a), upon which the edge of G2 would crop the beam as shown in Figure 2(c). We then adjust the axial position of L6 while compensating the change in beam path length between L6 and L7 using PS2. When the mirror edge of G2 is brought in focus in CAM2, G2 is aligned into the CBFP near L6 and can be set back to the *normal configuration* by adjusting the Y-axis on the G2-mount shown in Figure 2(a).

Next, we align the 4f system (L4, L5) in the *RGS* module. L4 and L5 are mounted in 16 mm cage mount (SCP05, Thorlabs Inc.) allowing for XY-translational adjustments, and are adapted to the 30 mm cage system supported by a shared pedestal post. This configuration allows the 4f system to be handled as an independent module. We first perform the relative positioning between L4 and L5 using a separate collimated laser beam outside of the LLSM, after which the distance and the relative in-plane lateral positions between L4 and L5 are finalized. The 4f system is then mounted into the LLSM beam path, and the adjustments of the lateral positions are performed by adjusting the lateral translation on the mounts for L4 and L5 concurrently and ensuring the excitation beam is centered at the optical axis with proper beam symmetry and position as inspected in both CAM1 and CAM2.

The next step is to fine-tune the position of G1. As shown in Figure 2(a), we set G1 into the *alignment configuration* by adjusting the X-axis in the G1-mount until the mirror edge of G1 is cropping the beam as shown in Figure 2(d). In *RGS*, the perpendicular-plane mirror (pp-M) and G1 are mounted on the same translational stage to ensure that the input and output beam positions either before pp-M or after G1 are independent from the axial position of G1. To position G1 into the conjugated back focal plan near L4, we adjust the Z-axis on the G1-mount until the mirror edge of G1 is in focus in CAM2, after which we can set G1 back to the *normal configuration*.

The *RGS* module along with the path-stretching modules decouple the degrees of freedom in the aligning process, and facilitate the precise positioning of G1 and G2 for both the initial construction of the microscope and the day-to-day maintenance. Additionally, in the *RGS* design, the G1 and G2 are mounted in rotational mounts, which is helpful for the maintenance when unexpected tilt is introduced from accidental galvanometer failure and crashing of the LLSM controller software, which, to our experience, has a chance to introduce a small tilt in the zero-voltage position for G1 and G2 and cause nonnegligible degradation in the light sheet quality. In that case, a simple fine adjustment on the rotational mounts used in *RGS* for the galvanometers allows us to fix the beam path without disturbing the setup.

The rest of the alignment procedures remain the same as the original LLSM design which can be obtained from HHMI through a no-cost license.

### 2.3. Operation configuration and maintenance

We built the LLSM in a BSL-1 lab space. To provide BSL-2 containment for imaging samples with *Legionella*, we used a custom designed enclosure (bioBUBBLE Inc), which is directedly mounted on the floor and isolated from the optical system shown in Figure 2(b) as the outer layer of enclosure with transparent side walls and a HEPA filter at the top. Our LLSM is also equipped with a custom-built laser enclosure that covers the entire top of the optical table using 20/80 rails and aluminum panels (G. A. Wirth Company, Inc) as shown in Figure 2(b) as the inner layer enclosure. The full-table laser enclosure blocks off the class-3 laser hazards, and also prevents dust drawn into the bioBUBBLE system from falling onto the optical elements.

The sample stage is maintained at 37 °C with humidity and CO2 control (Okolab). In our experience, the thermal drift induced with temperature fluctuations in the small range of 1 to 2 degrees inside the laser enclosure can reduce the light sheet quality and often necessitates manual adjustments of the alignment. Therefore, after constructing and aligning the microscope at room temperature, we kept the entire system operating non-stop (except for the lasers) with both the laser enclosure and the bioBUBBLE system closed, which can maintain the temperature gradient throughout the setup at a steady state comparable to the final operational status in addition to their primary function. We continued with daily fine adjustments with each session controlled below 30 minutes. The small adjustment time-window is critical in our application, because alignment requires opening up the enclosures which leads to temperature decreases that further induce noticeable reversion of thermal drifts in the system. In our experience, after approximately 2 weeks of such daily alignment, the instrument reaches a steady state without need for the manual optical alignment. In occasions when the whole setup cooled down, the alignment would be off at the beginning of a cold start, but warming up the instrument overnight solves the issue and can get the microscope ready for imaging without explicit manual adjustments on the optics. We achieved adequate instrument stability for imaging experiments spanned over 2 months, over which time we collected our datasets (discussed in section 4.2) without further manual optical alignment.

We checked the cell viability in an imaging experiment where a sample was imaged on LLSM up to 22.5 hours. As shown in Figure 3, we performed three time-lapse imaging sessions of the sample annotated as time blocks 1 to 3 to validate the effectiveness of the sample environmental control on the microscope stage. We observed multiple cell-splitting events in the early, middle, and late stages of the imaging session as shown in Figure 3(b), indicating that our instrument configuration is able to maintain sufficient cell health and the normal cell growth cycle. Time block 1 and time block 2 were acquired from the same field of view but separated by approximately 5 hours, each capturing cell-splitting events, indicating that the imaging process is gentle enough to keep the cell cycle robust. Time block 3 was acquired on a different area at the later phase of the imaging session, which also captured a cell splitting event. The full time-lapse movies are available in visualization 1. We also checked the photodamage of our final data acquisition routine (details are explained in section 3.4) where the sample is imaged non-stop for 50 time points. Here, as annotated in Figure 3(a), we imaged 3 groups of cells, and revisited the same region after 3 to 4 hours to confirm that the imaged cells were still alive and active (visualization 2), which validates that our acquisition routine is causing negligible photodamage to the cell.

**Fig. 3.**
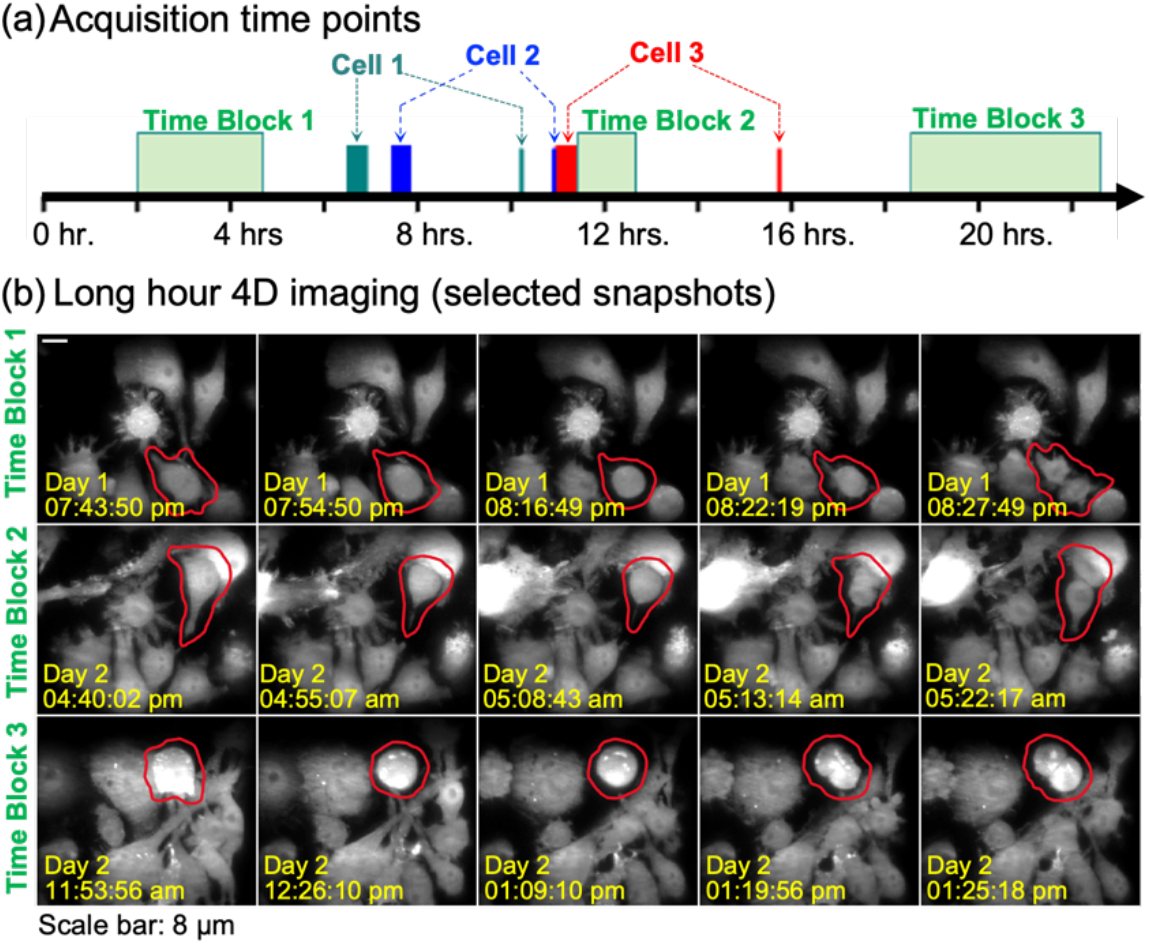
Cell viability validation. (a) shows the timeline of data acquisition on the same sample over 22.5 hours. The time blocks are marked on the time axis as rectangular regions with the edges matched to the start and end time of the data acquisition. Where block 1, block 2 and block 3 are time-lapse imaging blocks, and cell 1, cell 2 and cell 3 were imaged with the finalized data acquisition configuration (shown as the wider blocks) and revisited after approximately 3 to 5 hours to verify that the cell post imaging is still alive and active. (b) shows selective snapshots from the time series acquired for block 1, block 2 and block 3 featuring three cell splitting events that validate the cell viability on the microscope.

Last, we checked the stability of the system shown by imaging a fluorescence bead coated coverslips for 100 hours. The data acquisition is designed to mimic the actual data acquisition scheme to be discussed in section 3.2, where the *autofocus bead* step offered in the LLSM controller software is used to refresh the offset of the light sheet position, but only 1 volume of data were imaged per time point to reduce the total data volume for this characterization. The maximum intensity projections of the acquired volumes in XY, XZ and YZ planes over the entire 100 hours’ time course are shown in visualization 6, where we can see that there are minor drifts but autofocus bead was sufficient to maintain the alignment. The last frame we see a quick drawback of the imaging quality, that was caused by the water evaporation where the liquid level inside the sample chamber reduced below the tips of the objectives, causing aberrations to the light sheet. In principle, our modified design of the LLSM optical layout and entire instrument configuration should be able to offer the stability much longer than 100 hours and the imaging time can be further extended with the addition of a media perfusion unit. The characterization of the PSF and light sheet quality are shown in the Appendix Fig. A1.

### 2.4. Imaging specification

While the LLSM is designed with very rich flexibility for various of imaging configurations, here we focused on a configuration specifically designed for the study of *Legionella* infection on the macrophage cells. We chose the configuration optimized over phototoxicity and photobleaching, imaging duration, availability of biologically meaningful information, the size of the 3D field of view to accommodate cell movement, and attainability of the overall workflow within the timescale of the experiment. We used the annular mask with inner and outer numerical apertures of 0.35 and 0.4 respectively, and calculated the SLM pattern to obtain a light sheet with a theoretical 30 μm FWHM based on the optimization strategy outlined by Chen, et al [18]. We use the objective scanning mode to image each time point for 301 slices spaced with 100 nm intervals, with approximately 100 ms exposure time per slice. The final acquisition is approximately 30 seconds per 100 μm × 50 μm × 30 μm volume with focal volume of roughly 100 μm × 30 μm × 30 μm. For each cell, 50 time points are collected in the red channel only to capture the cell dynamics. When imaging the infected macrophage cells, we collect 1 time point in the green channel to confirm that the rod-shape structure observed in the red channel are indeed *Legionella* prior to the 50 time points acquisition in the red channel.

Figure 4 demonstrate the typical imaging result of a macrophage cell infected with *Legionella* using our imaging routine. The green channel (Figure 4(a)) captures the *Legionella* labeled with CellBrite^®^ Fix 488 and the collagen labeled with FITC, each displayed with the optical pixel intensity dynamic range to focus on *Legionella* (left) or *collagen* (right). The fluorescent collagen fibers guide us to mount the sample effectively, serve as internal marks to confirm the imaging quality for each specific dataset, and confirms that the microenvironment experienced by each cell are similar as gauged by the size and density of the collagen fibers, and ensure we are not imaging cells embedded in a region where the collagen is thicker or thinner due to the inhomogeneity in the polymerization process for the collagen. The dye labeled *Legionella* help us to confirm that the rod-shape feature observed are indeed *Legionella*. The red channel (Figure 4(a), middle) captures the macrophage cell and *Legionella* both engineered to have cytosolic mCherry expression. Note that not all the *Legionella* cells have mCherry expression in the final imaging condition, as shown in Figure 4(a) where the intersect misses a grain-shape feature when compared to the left panel (red arrow). The imperfection of mCherry expression in *Legionella* does not influence the results of this study but can be further improved with genome integration of the mCherry sequence in *Legionella*. We segmented the peripheral and internal region of the cell through data processing and display the two parts with different color schemes as shown in Figure 4(b). Details of the sample preparation and data analysis are explained in the following sections.

**Fig. 4.**
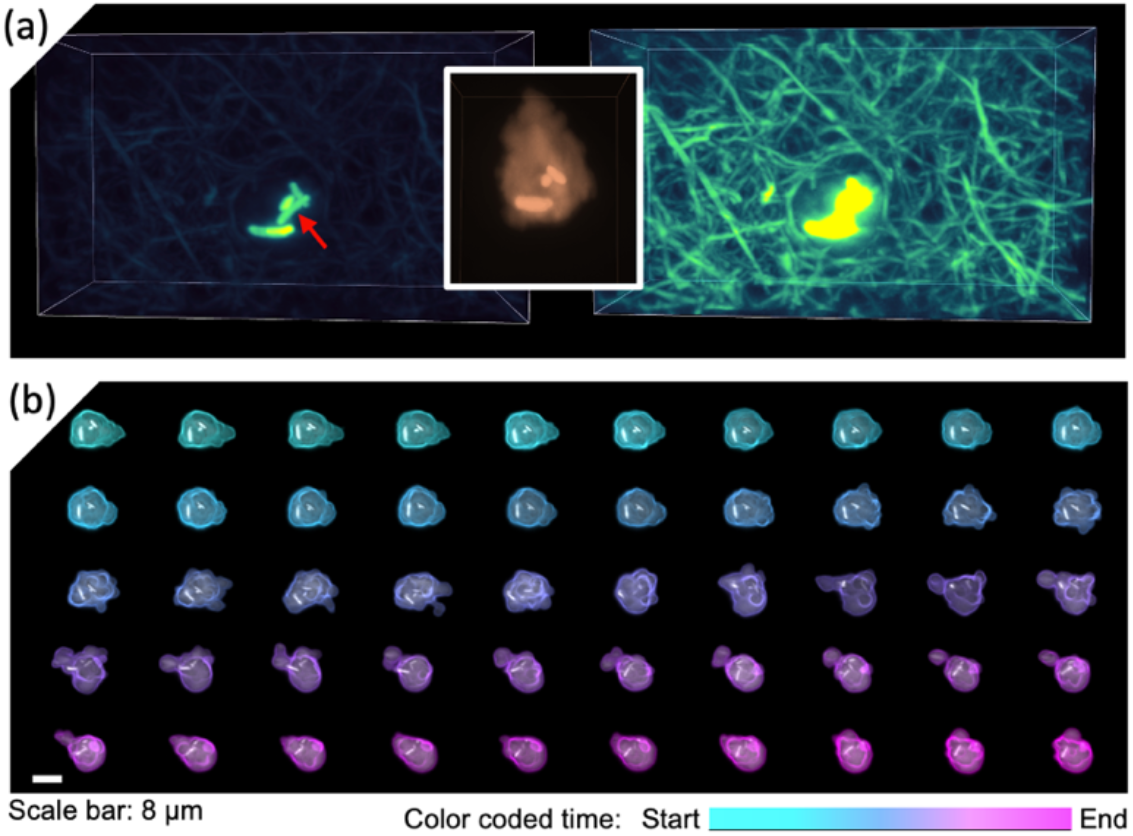
Imaging result of one cell. (a) Shows the 3D rendered (with maximum intensity projection) imaging result in the green channel displayed with the dynamic ranges focused on *Legionella* with CellBrite^®^ Fix 488 stain (left) and FITC-labeled collagen matrix (right). The red channel captures both the macrophage cell and the Legionella with mCherry emission (intersect). The red arrow indicates a *Legioenlla* bacteria without mCherry expression. (b) visualization of the 50 timepoints of the cell imaged over 26 minutes. The cell peripheral and internal regions are separated and displayed independently, where the cell peripheral region is color coded in time, and the cell internal region is displayed at gray scale. The grain-shape feature highlights the position of the *Legionella*.

## 3. Experiment

We designed the experiments to study the *Legionella* infection on macrophage. The *Legionella* strain used in this study is modified from the DotF-sfGFP (JV9082) strain [21] obtained from Grant Jensen’s group as a gift. We used electroporation to transform the mCherry encoding plasmid pON.mCherry [22] in the JV9082 strain and selected the red colonies as the initial strain to support our experiments. The pON.mCherry was a gift from Howard Shuman (Addgene plasmid # 84821). In our study, we found that the plasmid was preserved well with antibiotic selection pressure, and the mCherry expression is adequately maintained without selection under the conditions of our experiments. Note that the mCherry expression level tends to reduce when the selection pressure is relieved; therefore, we limited the usage of our *Legionella* without the selection pressure to less than 24 hours. For the macrophage cells, we obtained RAW264.7 cell line from ATCC and used retroviral transduction to engineer a RAW264.7 cell line with constitutive, cytosolic expression of mCherry. RAW264.7 is a murine macrophage-like cell line commonly used as a model system for macrophage cells. For simplicity, in the rest of this manuscript, we use the general term *Legionella* to refer to our engineered mCherry expressing *Legionella pneumophila* strain, and we use the general term *macrophage* to refer to our engineered RAW264.7 cells with mCherry expression. Although we engineered both the *Legionella* and the macrophage cells for cytosolic mCherry expression, the mCherry concentration inside the *Legionella* is higher than that in the macrophage cells, which provides sufficient contrast for us to distinguish *Legionella* from the macrophage cell cytosol. The *Legionella* cell is stained with CellBrite^®^ Fix 488 stain (Biotium, Cat. No. 30090) and imaged for one time point in the green channel before the 4D imaging time series to confirm that the grain-shape structures with higher mCherry signal level inside the macrophage cells are indeed *Legionella*. For the 4D time series data acquisition, we used the same color channel to image both species to reduce both the time cost and dosage of light exposed to the sample, which favors both time resolution and signal level. Details of the experiments are explained in the following subsections.

### 3.1. Sample preparation

We prepared samples under two different conditions where the macrophage cells are prepared either without or with *Legionella* infection, and will be addressed as “naïve macrophage” and infected “infected macrophage” respectively. As shown in Figure 5, the processes can be grouped into two categories: first, the preparatory tasks (highlighted with green boxes, Figure 5(a)(d)(e)) that are performed as continuous processes on a relatively flexible experimental schedule, and second, the on-demand tasks (highlighted with gray boxes, Figure 5(b)(c)(f)(g)(h)) that are performed on the day of imaging experiment right before each imaging session.

**Fig. 5.**
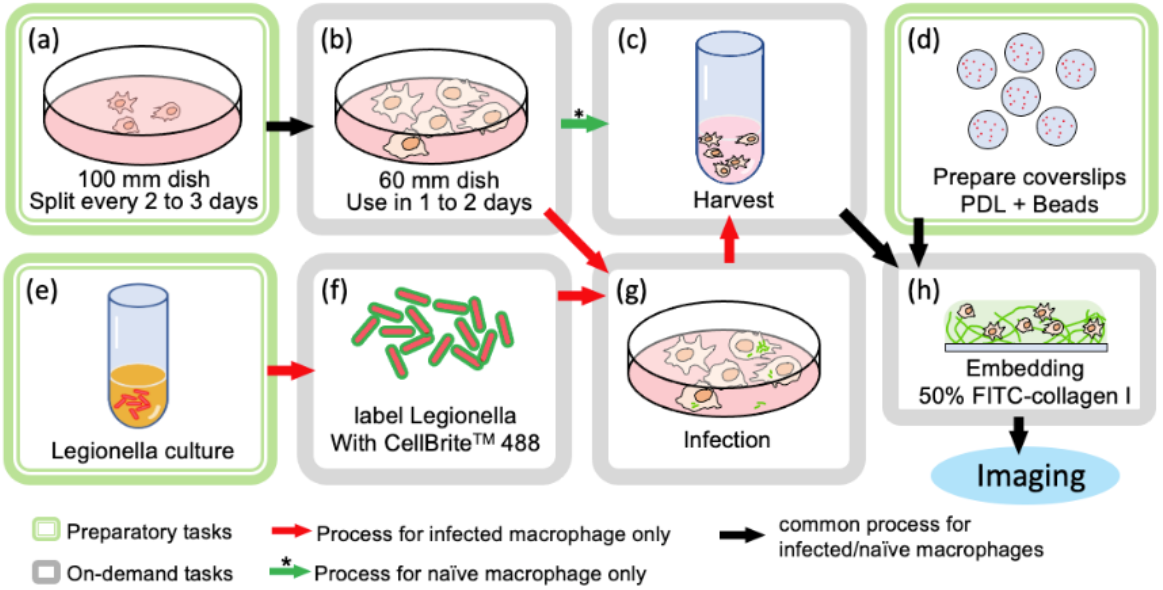
Sample preparation workflow. The sample preparation tasks can be grouped into preparatory tasks (a)(d)(e) and on-demand tasks (b)(c)(f)(g)(h). The sequences of the tasks are indicated with the arrows.

The preparatory tasks include the culturing of macrophage cells (Figure 5(a)), preparation of imaging coverslips (Figure 5(d)), and the culturing of *Legionella* (Figure 5(e)). To maintain the supply of macrophage cells for the experiments (Figure 5(a)), we kept an active line of the cells cultured in 100 mm cell culture plate using DMEM (ThermoFisher, Cat. No. 10313039) supplemented with 10% fetal bovine serum (FBS), 1× Penicillin-Streptomycin-Glutamine (ThermoFisher, Cat. No. 10378016), 2 mM L-glutamine (fisherscientific, Gibco™ 25030081) and 4 ug/ml puromycin. The cells were used up to the 20^th^ passage until replaced by defrosting a new cryogenic stock from the stocks frozen from the same batch. The cells were used for experiments after two passages post defrosting. We passed the macrophage cells by loosening the cell adhesion using 0.05% trypsin applied for minimum of 10 minutes followed by flushing the cell culture media directly onto the cells using a 1 ml pipet. When cells cannot be lifted, we incubated the cells with 0.05% trypsin for an addition of 10 minutes before lifting with the media flushing method. The cells were then pelleted through centrifuge at 150×g for 5 minutes, counted, and seeded in 100 mm plate with fresh complete media. For each passage, we seeded either 1.2×10^7^ or 6×10^6^ cells for the next passage in 2 or 3 days, respectively. To prepare the imaging coverslip (Figure 5(d)), we first cleaned the coverslip with 70% ethanol, wiped with a clean Kim wipe followed with an air-dry step inside an operating biosafety cabinet for 10 to 30 minutes, then we coated the coverslip with Poly-D-lysine (Sigma-Aldrich, CAS No. 27964-99-4) following the manufacture’s protocol. We further coated the PDL treated coverslip with fluorescent beads (ThermoFisher, FluoSpheres™ Carboxylate-Modified Microspheres) to serve as in-situ fiduciary markers for autofocus during an imaging session. The coverslips are stored at room temperature in dark and used for imaging within a month. To inoculate the *Legionella* strain (Figure 5(e)), we streak the strain from a frozen glycerol stock on a CYE plate supplemented with thymidine (0.1 mg/mL) and chloramphenicol (5 ug/mL) followed with growth at 37 °C for 3-4 days. A colony was inoculated in AYE media supplemented with thymidine and chloramphenicol and further grown for 20 hours.

The on-demand tasks (Figure 5(b)(c)(f)(g)) are performed in accordance with the imaging experiment. It contains two paths that provide samples of either naive or infected macrophage cells. As shown in Figure 5, both processes start from a portion of macrophage cells seeded in 60 mm cell culture plate (Figure 5(b)) prepared 1 or 2 days before the imaging experiment in parallel with the corresponding cell passaging experiments (Figure 5(a)), from which 6×10^6^ or 3×10^6^ cells are plated in the 60 mm dish for preparing imaging samples in the following 1 or 2 days, respectively.

To prepare the infected samples, we prepare the *Legionella* bacteria (Figure 5(f)) one day before the imaging experiment using a series of inoculation tubes containing initial OD600 values ranging from 0.1 to 0.3, and grown at 37°C in cell shaker for 20 hours. We measure the OD600 of the culture inoculation upon harvest and choose the tube with OD value closest to 3 to proceed. The bacteria are then pelleted and resuspended in 1× PBS buffer with OD600 adjusted to 1 to 1.5, from which the bacterial suspension is stained with CellBrite Fix 488 (Biotium, Cat. No. 30090-T) following the manufacturer’s protocol using 5× of the stains, and adjusted to the original volume with OD600 at 1 using DMEM without phenol red (ThermoFisher, Cat. No. 31053036). Next, we used the stained *Legionella* to infect the macrophage cells (Figure 5(g)) by removing all the cell culture media from the 60 mm dish, apply 0.5 ml of the stained *Legionella* suspended in DMEM, centrifuge at 150 × g for 10 minutes to synchronize the infection, and incubated inside the cell culture incubator for 1 hour to allow for the initiation of phagocytosis uptake of *Legionella* by the macrophage cells. Then the cell dish is gently washed to remove the excess amount of *Legionella* and the media is replaced with complete DMEM without phenol red supplemented with 10 ug/ml gentamycin to kill the extracellular *Legionalla*. After one additional hour of incubation that allows for full internalization of *Legionella*, the cells are ready for the harvesting step (Figure 5(c)). When working with the naïve macrophage condition, we directly proceed from the dish of cells in 60 mm cell culture plate without infection.

To harvest the cells from the cell culture dish (Figure 5(c)), we rinse the cells twice each with 5 ml of 1× PBS buffer, and apply 3 ml of 0.05% trypsin followed with 10 minutes of incubation to loosen the cell adhesion, then lift the cells by flushing the media directly on the cells using a 1 ml pipet. The cells are then pelleted by centrifuging at 150 × g for 5 minutes, counted and resuspended in DMEM with final concentration adjusted into the range of 6×10^6^ to 1.2×10^7^ cells per ml.

The next step is to embed the harvested macrophage cells in collagen matrix directly on top of the imaging coverslip with PDL coating and fluorescence beads (Figure 5(h)). We first prepare the neutralized collagen mixture which is a 2 mg/ml collagen solution suspended in 1× PBS with 1:1 ratio of unlabeled Type-I collagen (ThermoFisher, Cat. No. A1048301) and FITC labeled Type-I collagen (Sigma-Aldrich, Cat. No. C4361-10ML). Because the collagen starts to polymerize as soon as the mixture is neutralized, we prepared aliquots of buffer and aliquots of pre-mixed collagens stored at 4°C prior to use, and directly mix the two aliquots right after the harvest step (Figure 5(h)). We combine neutralized collagen mixture and the cell suspension at 1:1 ratio, and transfer 15 μl of the cell-collagen mixture to the coverslip, allow the liquid to access all surface of the coverslip, then remove 12.5 μl of the mixture from the coverslip. We then incubate the sample in a 37°C cell culture incubator with 5% CO2 for 20 minutes for collagen to polymerize, the residue forms into a collagen pad that embeds the cells for imaging under the LLSM. The theoretical collagen pad thickness is 127 micrometers assuming the pad is perfectly flat. In practice, a curved top surface is obtained due to surface tension, and the collagen pad thickness is in the range of 100 μm to 200 μm. This embedding step is performed inside a 6-well cell culture dish, and we finish this step by adding additional 2ml of DMEM without phenol red to the sample to fully immerse the collagen pad and incubate the cells in the CO2 cell culture incubator for 30 minutes before mounting and imaging on LLSM. Note that the cells are turbid medium which can cause distortions to light sheet quality and aberrations to the imaging result. Our protocol provides approximately 7500 to 15000 cells in the sample, the estimated lateral distances between the projected positions of the cells at the coverslip surface would be 37.4 μm to 51.2 μm, and the estimated average distance between cells in the 3D matrix is 58.6 μm to 66.9 μm assuming the cells occupy regular grids. Therefore, the cells are well separated such that we can effectively choose to image the cells without adjacent cells that impacts the imaging quality. Also, the polymerized collagen fibers are thin and fibrous structures (shown in Fig. 4(a)) that do not create piece wide constant distribution of different refractive indexes across the 3D environment, therefore the aberrations caused by the embedding gel in both the excitation and detection light field are minimum.

In summary, our sample preparation procedure is designed to provide an attainable workflow for small research teams when imaging and data acquisition processes are involved concurrently (discussed in section 3.4 and 4.2). One imaging session can be finished in one day with the flexibility to be extended longer. All the preparatory tasks can be maintained with 2 to 3 days of intermittence. On a day of imaging experiments, we perform the on-demand preparation steps in the morning and perform data acquisition for the rest of the day. In our experience, when working with naïve macrophage cells, all the sample preparation work can be finished in the morning. When working with infected macrophage cells, the on-demand tasks can be achieved within 5 hours, after which the cells are mounted on LLSM ready for imaging. The post-infection time is approximately 3 hours when the cells are ready for imaging. When imaging the infected macrophage cells, we control the imaging session to be 5 hours, therefore all the cells are imaged within a post-infection time window of 3 to 8 hours, corresponding to the intra-cellular growth phase of the internalized *Legionella* as discussed in [23], ensuring that the acquired dataset is representing a relatively monotonic phase over the entire time course of the relevant host-pathogen interaction events.

After the protocols are established, we are able to maintain the workflow with efforts from 1 to 2 researchers. Specifically, for the condition with naïve macrophage cells, we performed our experiments solely by 1 researcher. For condition with infected macrophage cells, 1 researcher is sufficient to prepare and image one sample on an imaging day, and with 2 hours of support from an additional researcher we have attained the throughput to prepare and image 2 samples per day. The efficiency can be further improved with better automated data acquisition routines (discussed in section 4.2), and we believe our workflow is also transferrable for better staffed and equipped imaging centers to facilitate effective collaborations and services.

### 3.2. Data acquisition routine

We imaged individual macrophage cells (either naïve or infected) continuously for approximately 26 minutes to obtain 50 time points of observation for each cell. Note that the total time contains small variations in the milliseconds range originating from resetting the instrument between consecutive imaging time points such as stage movement and stabilization, as well as resetting the camera sensors. Outside of an imaging experiment when the instrument was kept at idle state, all parts except for the lasers were powered on, and both the laser enclosure and the bioBUBBLE enclosure were kept closed to maintain the thermal gradient across the instrument in a status close to that of an imaging session. Inside an imaging session, our data acquisition workflow is shown in Figure 6.

**Fig. 6.**
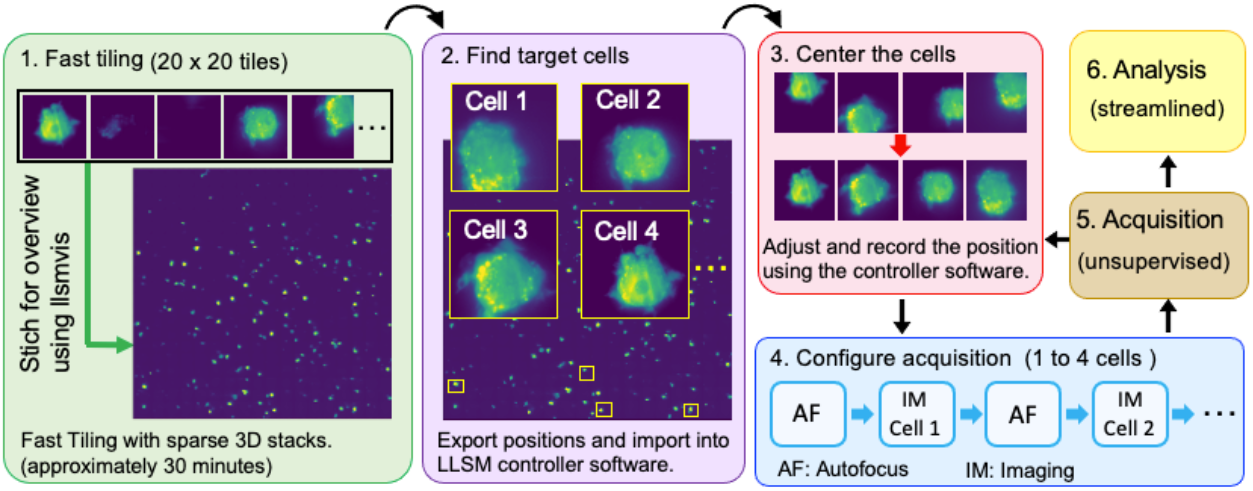
Data acquisition workflow. The data acquisition workflow is composed of a series of preconfigured imaging routines, data processing steps and manual data acquisition control. Step 1 involves manual sample mounting and acquisition with pre-configured fast tilling routine, and all the data is processed on a cluster on demand. Step 2 involves manual selection of the cell of interest. Step 3 is a manual step to position the cell to the center of the focal volume, for which further automation is plausible. Step 4 is a semi-automated process, where the acquisition routine is pre-configured, and manual configuration only involves updating the position coordinates of the fluorescence beads and cells. Step 5 is an unsupervised data acquisition step after which we either move on to further data analysis or continue with the data acquisition by looping back to Step 3.

To start an imaging session, we first wipe the sample stage with 70% ethanol and let the residual ethanol evaporate completely for approximately 30 minutes. We then we apply 7 milliliters of prewarmed DMEM without phenol red into the sample holder bath on the LLSM and allow the system to settle to equilibrium for at approximately 30 minutes with the two layers of enclosures closed. We use this waiting period to mount the imaging sample coverslip into the sample holder and mount the holder to the microscope. The sample should be mounted onto the microscope when the sample stage has achieved stabilized temperature at 37°C as indicated by the microscope top chamber system (Okolab). Carefully position the sample and adjust the mounting screws until the collagen is visible within field of view through the imaging camera (CAM3 as shown in Figure 1) when the sample stage is set at a pre-configured height for the fast-tiling step shown in Figure 6. Let the sample sit in the microscope with the two layers of enclosures closed for at least 30 minutes to allow the temperature gradient as well as the micro drifts introduced during the sample mounting step to settle. We use this time window to perform Step 4, where we navigate in the sample to look for several internal fiduciary markers that will be used for auto-focus, and configure the data acquisition routine.

We then perform fast tiling over a large area with sparse sampling steps as shown in Figure 6, Step 1. This acquisition routine should be preconfigured in the controller software to save time and the preparation step here should only involve emptying the pre-configured data folders. The results are transferred to a separate PC such that the data can be managed completely independently from the instrument controlling PC. In our workflow, we uploaded the data to the high-performance computing facility (Livermore Computing) for massive processing on demand. The results of the tiled scan are accessed through a Jupyter notebook to allow for user inspection, from which we identify the indices of the tiles that enclose cells as shown in Figure 6, Step 2. From these tile indices, our processing routine generate a .csv file enclosing the location coordinates of the tiles, which can be imported into the LLSM controller software for the follow-up acquisition configurations.

When we are ready to image the cells, we navigate to the positions of the selected tile that captured the cell, and manually refine the position of a cell to position it to the center of the focal volume as shown in Figure 6, Step 3. We image the cells in batches each with 1 to 4 cells, because after the cell coordinates are configured, the cell within the same batch still has a time window to migrate before it gets imaged. In our experience, controlling the batch size under 4 (time window under 2 hours) is sufficient to keep majority of the imaged cells within the volume of interest. The larger the batch size, the less effort is required from the operator. We acknowledge that an automated routine to detect and set cell position would be an elegant solution to compensate for the cell movement and further reduce the effort required from the researcher, but that is beyond the scope of this study. The imaging acquisition routine for each cell includes four steps as shown in Figure 6, Step 4: visit the position of a fiduciary marker, perform the auto-focus procedure, revert to the position of the cell, and image the cell. We configure the process using the scripting tool available in the LLSM controller software with minimum interference from the researcher. At this stage on an imaging day, the instrument operation required from the researcher involves only preparing and mounting the sample, and one sample supports imaging sessions of many hours. In this study, the imaging session lasts for up to 2 days and is primarily limited by the cell health inside the microscope or the biological relevant time course. Longer imaging sessions are plausible by integrating a perfusion system to refresh the cell culture media on the microscope. Therefore, the workflow is well suited for remote instrument control which can greatly improve the work efficiency and throughput. Our remote control is easily facilitated by the Microsoft Remote Desktop tool that offers remote access to the instrument controlling computer.

Using this data acquisition procedure, we acquired 4D datasets of 268 cells, each for approximately 26 minutes over 20 days of imaging experiments. For the condition with naïve macrophage cells, we imaged 157 cells from 10 samples over 10 imaging sessions, one imaging session lasting for 1 to 2 days. For the infected macrophage cells, we imaged 111 cells from 15 samples over 15 imaging sessions, one session lasting for up to 5 hours to focus on the intracellular bacterial replication phase of Legionella infection. Among all the datasets acquired using this routine, 13.1% are identified as outliers for reasons including interference with adjacent cells and the associated difficulty for segmentation, cells moving outside of the field of view, cell death and several rare cellular events that are beyond the scope of this study (e.g. cell splitting, macrophage clearance [24].). The imaging experiment is designed to provide gentle and continuous live cell imaging of cells embedded in collagen pad of 100 to 200 microns thick, which is a type of observation uniquely provided by LLSM as compared to the commonly accessible commercial microscopes. Our imaging workflow provides the throughput and efficiency that accommodates the uncertainties needed in the follow-up studies that often require multiple probing conditions with carefully designed molecular biology experiments, which is beyond the scope of this study but well suited as a follow-up study. The process is semi-automated with minimum interference from the researcher; therefore, it is attainable for small research teams. We also believe further adoption of this imaging workflow by the imaging centers and the relevant commercial product is plausible and beneficial for the broader research community to promote the application of LLSM.

## 4. Data analysis

We developed custom-designed analysis methods to analyze the biologically relevant information from the acquired datasets. On one hand, it contains the general functions for the coarse processing of LLSM datasets including parsing and organizing the datasets all saved in a single folder, deskewing, generating .MP4 movies of maximum intensity projections of the datasets, generating data acquisition parameter reports, etc. On the other hand, it also contains the tailored analysis methods designed specifically in this work. We acknowledge that more comprehensive morphology analysis can be performed using prior methods such as u-shape3D [25] but is beyond the scope of this work. Our analysis scripts are currently open source and packaged as the *llsmvis* repository on GitHub with a collection of Jupyter Notebook examples to facilitate the community adoption of our workflow [32]. We performed our analysis partially on a desktop computer and mostly on the high-performance computing (HPC) facility provided by Livermore Computing. Our scripts are purely developed in Python and are transferrable for publicly available HPC facilities such as XSEDE and Amazon AWS; it can also be configured on independent computers in scenarios when the increased data processing time is tolerable. Details of the data processing pipeline and the tailored analysis methods are explained below.

### 4.1. Data processing pipeline

The analysis pipeline is shown in Figure 7. Each output dataset placed by the LLSM controller software is a collection of TIFF image stacks *I*(*t*) and one *Settings.txt file that specifies the data acquisition configuration information for the stack. In our imaging experiments, we save all the datafiles to the same folder without intensive organization effort, allowing the researcher to focus most of the attention on imaging the sample. When processing the data, as indicated in Figure 7(a), our analysis routine first detect the dataset information files (*Settings.txt files), distinguish and parse the information for each dataset; and process the TIFF stacks into ordinary 3D stacks (i.e. the deskew step), and the maximum intensity projections (MIP) are calculated for all the image stacks in XY, XZ and YZ planes to obtain the MIP image series M_xy_(t), M_xz_(t) and M_yz_(t) respectively. We then calculate the time-integrated MIP images for each plane to capture the overall range of the cell inside the volume of interest. Next, as shown in Figure 7(b), we wrote an interactive cropping tool to mark the region-of-interest from the time-integrated MIP images in all three planes, from which we generate a new series of volumes that contain only the signal from the user-defined volume-of-interest (Figure 7(c)), which is zero-padded into tight-bounded 3D volumes and stored as TIFF stacks (the trimmed stacks).

**Fig. 7.**
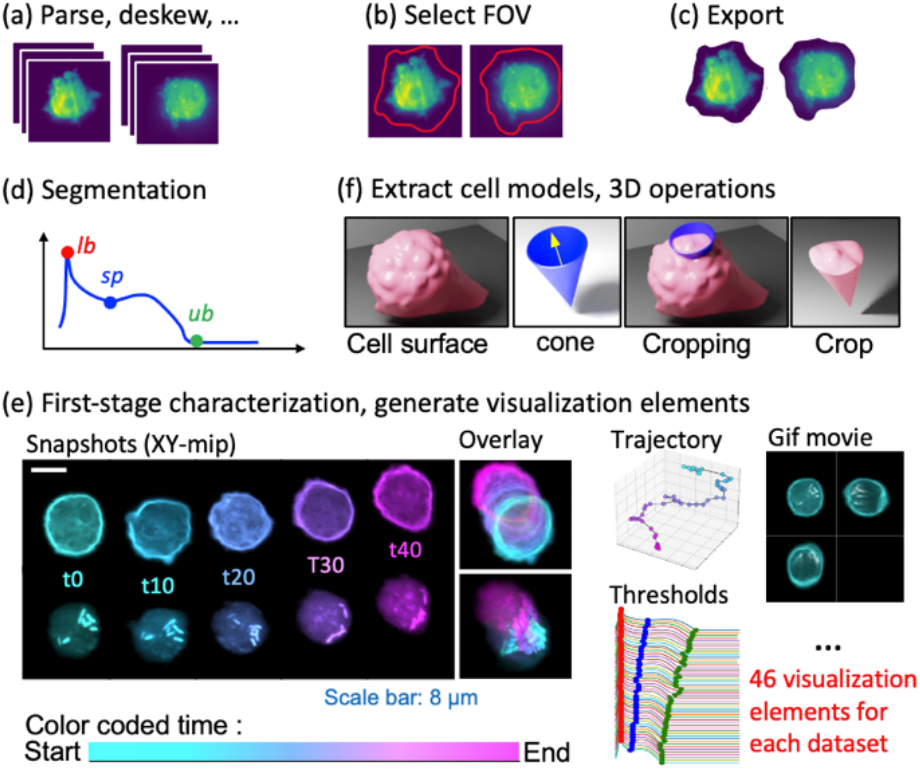
Data processing routine. (a) performs the basic transforms organizations of the raw data for the follow up analysis. (b) involves an interactive tool that allows the user to manually select the cell of interest to process. (c) export the selected region of interest into independent TIFF stacks which can reduces the size of the dataset and reduce the subsequenct computational cost. (d) is an automatic step that generates a characterization profile of the voxel intensity distributions (indicated by blue curve) to segment the cell volume into cell peripheral and internal region using three signature points indicated as the *lb*, *sp* and *ub* points. (e) Is a streamlined analysis step that performs the characterization analysis, and also generate 46 visualization elements for each dataset to facilitate user inspection. (f) extract the cell models and perform the follow up 3D operations to support further analysis to be discussed in section 5.2 to 5.3. The data analysis routine can either be processed in batch over all datasets, or be performed on individual datasets. All the analysis scripts are open source on our GitHub repository *llsmvis* [32].

The fourth step (Figure 7(d)) is to segment the voxels in each 3D volume into background, cell surface and cell internal regions. Specifically, we calculate the probability histogram of the voxel intensities denoted as *h*(*i*), where *i* represents the voxel intensity. Next, we transform *h*(*i*) into a characteristic profile *C*(*i*)=(*h*(*i*)+1)^0.01^ shown in Figure 7(d). The principles for the design of *C*(*i*) are that *C*(*i*) should be easy to compute, *C*(*i*) should have similar profile across all time points across all datasets, and most importantly *C*(*i*) should display prominent signature points that represents the voxel intensity thresholds to be used for segmentation, and the signature points should be robustly identified through detection of local extremals of either *C*(*i*) or the different orders of derivative of *C*(*i*) when combined with universally defined smoothing criteria, manual adjustment is allowed but shouldn’t require too much effort. Three signature points on *C*(*i*) are used as shown in Figure 7(d), the lower bound (*lb*), the saddle point (*sp*) and the upper bound (*ub*). For a voxel with intensity *i*, when *i*<*sp*, the voxel is recognized as the background noise; when *sp*<*i*<*ub*, the voxel is recognized as the cell body; when *i*>*ub*, the voxel is recognized as an outlier voxel that is overly bright. The *lb* point represents the most abundant value in the background noise region, and is used as the lower bound for the pixel intensity dynamic range when displaying the image as a picture for the automated generation of the visualization elements, and the *ub* point is used as the upper bound of the dynamic range. The *sp* point is used to identify the cell peripheral region constitutes of the voxels with intensities below and up to the *sp* point, for which the total number of cell peripheral voxels are set to be a certain fraction of the total number of voxels within the cell body, in this study, we fix the fraction as 50%.

After the segmentation, a series of analyses are performed to characterize the cell morphology and the relevant dynamics as shown in Figure 7(e). The analyses include visualization of the cell body, internal region and peripheral region, and generate the corresponding .mp4 files and .gif files for users to inspection the dynamics. We also identified the geometry center of the cell at each time point and plot the corresponding cell migration trajectories, provide plot of the characteristic profile C(*i*) for user to inspect the robustness of characteristic points for segmentation, and allow for necessary manual adjustment, etc. Total of 46 visualization elements are generated and grouped in a folder designated for each cell for users to inspect. The visualization elements are organized in HTML pages to facilitate efficient browsing of the results, which are released on figshare [31] with the associated instructions released on the *llsmvis* repository on GitHub [32].

The next step (Figure 7(f)) is to extract the cell surface and perform the relevant operations and analyses using the resultant 3D cell model. We implemented our operations using the Python package of the Visualization Tool Kit (VTK). At this point, our data can be perceived as a three-dimensional scalar field with elements corresponding to voxels specified with location coordinates and intensities. We use the intensities to identify the voxels that belong to the cell body and achieve a unique description of the cell surface. Specifically, we use the *sp* point as a threshold to transform the scalar field into a binary mask, where voxel values 1 correspond to the cell body, and 0 to the background. Next, we apply a morphological opening and closing operation on the binary mask with a 3×3×3 kernel (with all elements equal to 1) to remove the imperfections on the binary mask as influenced by the noise, and extract the isosurface from the binary mask using the Marching Cubes algorithm [26]. A polygonal mesh is then extracted from the binary mask volume to represent the cell surface defined as the boundary between the cell body and the background. To ensure that we extract a tightly bounded cell body, we apply connected component analysis and keep only the largest connected component as a close representation of the cell surface. In the final step of surface extraction, we relax the point coordinates in the surface using Laplacian smoothing. The extracted cell surface is exported into a *.stl file for each time point, which can be loaded by VTK for the following up analysis, or imported into computer graphics software toolset such as Blender and Houdini for visualization. We also provide the tools to generate 3D animations of the dataset using Houdini under the no-cost apprentice license. As demonstrations, visualization 3 and visualization 4 demonstrate the representative videos for a naïve macrophage and an infected macrophage respectively.

Based on the extracted cell surface, we derive additional analysis steps to characterize (a) whole-cell smoothness, (b) patch-wise smoothness on local regions of the cell surface and (c) cell polarity. These three steps are expansions of the analysis step shown in Figure 7(f), and are discussed in detail below.

### 4.2. Whole-cell smoothness characterization

To compute the whole-cell smoothness as shown in Figure 8(a), we start from the cell surface, extract the cell volume (*V*) as the total volume enclosed by the cell surface, and extract the cell surface area (*A*). We then calculate the whole-cell smoothness as *S* = *R*_V_/*R*_A_, where *R*_V_ and *R*_A_ are the radiuses of spheres that share either the same cell volume (*V*), or the same cell surface area (*A*) with the 3D cell model. We acknowledge a perfect sphere as a perfectly smooth cell, which yields *S*=1. When any protrusion or recession structures are created on the perfect sphere, we will have *S*<1, indicating a decrease in the smoothness. We crafted this cell smoothness measure, *S*, as a dimensionless quantity that is independent of any length scale and bounded between 0 to 1.

**Fig. 8.**
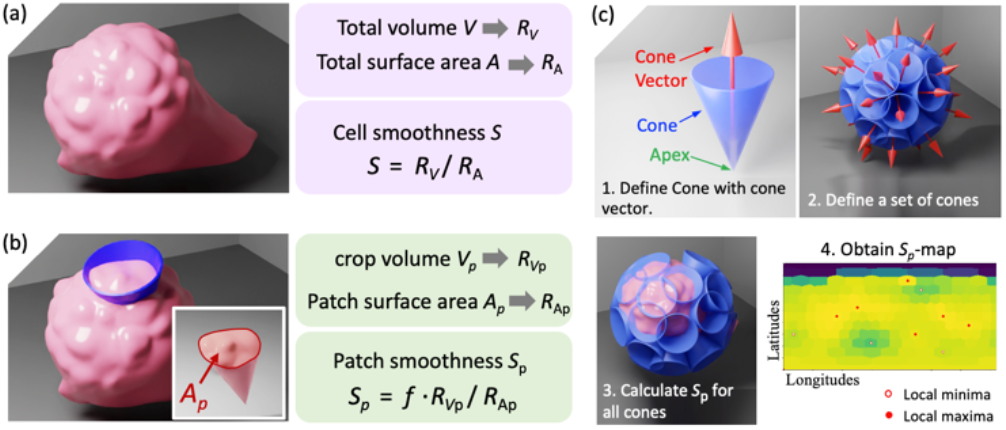
Cell smoothness characterization. (a) shows the process to characterize the whole cell smoothness. Left panel shows a 3D rendering of a cell surface and the right panels shows the methods to calculate cell smoothness *S*. (b) shows the process to characterize the patch-wise cell smoothness where a cone is used to crop the cell volume (left panel) to obtain a cropped-out volume and a cropped-out cell surface patch (intersect of left panel), and are used to deduce the patch-wise smoothness *S*_p_ (right panels). (c) shows how we extend the patch-wise smoothness characterization across all directions for the cell by defining a set of cones, calculating *S*_p_ for each cone and generate a *S*_p_-map for visual inspection.

### 4.3. Patch-wise smoothness characterization

Similarly, the patch-wise smoothness is computed based on *patches* of the cell surface as shown in Figure 8(b). We first define a cone *Cone*(***r***_c_, α) as shown in Figure 8(c), where we define ***r***_a_ as the cone vector, which is a unit vector pointing from the cone-apex towards the center of the cone-base to specify the cone orientation, and a is the apex angle of the cone. The apex of the cone is placed at the centroid of the cell volume defined as the volume enclosed by the cell surface discussed in section 4.1, and the height of the cone should be sufficient to allow the cone side surface to penetrate through the cell surface as indicated in Figure 8(b). In our case, we set the cone height to be the longest edge of the focal volume to ensure all cones penetrate through the cell surface. We take the intersection between the side surface of the cone (*F*_cone_) and the cell surface (*F*_cell_), and obtain the boundary curve *B*_cc_ = *F*_cone_ ∩ *F*_cell_ (indicated by the red line shown in Figure 8(b) intersect). For each cone, the *patch* of interest is defined as the patch on the cell surface *F*_cell_ cropped out by the cone and is bounded by the boundary curve *B*_cc_. In case of ambiguity where the cone crops out multiple patches from the cell surface, we choose the patch closest to the cone apex to proceed. This way, we ensure that when an elongated surface protrusion extends outside of the cone frustrum, the extended part will not be cut out and the patch includes the entire protrusion. We calculate the volume enclosed by the patch and the side surface of the cone inside the cell, and denote the volume as *V*_p_. We then obtain the total area of the *patch* (*A*_p_) as shown in Figure 8(b), and calculate the patch-wise smoothness value as *S_p_* = *f R*_Vp_/*R_Ap_*, where *f* is a normalization factor, and *R_Vp_* and *R_Ap_* are the radiuses of perfect spheres if the cone is cropping a perfect sphere and yields equivalent *A*_p_ and *V*_p_ respectively.

We then characterized the patch-wise smoothness around all directions of the cell at each time point from the 4D live cell imaging datasets by repeating such analysis over a collection of cones as shown in Figure 8(c). We fix the apex angle of the cones at 40° and use purely the cone vector to define each cone. Then we define a set of cones by defining a set of unit cone vectors with the vector heads located on a unit sphere and semi-equally spaced in distance, as illustrated in Figure 8(c). Specifically, we hold the angle between two adjacent vectors close to parameter ϕ, which is used to control the sampling rate of *S*_p_ in all directions and can be optimized for different feature of interest. In this study, we fixed ϕ=15°). Note that in the illustration shown in Figure 8(c) panel 2, the density of the cones is lower than the actual density used in the study, the purpose is to keep the illustration clear and perceivable demonstrating both overlapping and non-overlapping cones. Additionally, the cone apex angle is fixed at 40°, which is greater than ϕ, so in reality we have densely intersected cones to provide a sliding window averaging effect in the analysis results. We then calculate the *S*_p_ over all cones for the cell as illustrated in Figure 8(c) panel 3. Because here the cone vector is the only variable that defines the patch of cell surface cropped out by a cone (other variables such as the cell surface and the apex angle of the cone are fixed), so the path-wise smoothness quantity *S*_p_ can be expressed as a function on the polar coordinate system *S_p_*(θ, φ) where θ and φ are the angular components of the spherical coordinates of the cone vectors. Therefore, we calculated the *S_p_* defined by the cone set, which is equivalent to sampling the function *S_p_*(θ, φ) on a discrete support represented by the cone set. Next we characterized the local extremals of *S_p_*(θ, φ) by identifying the sampling points that has all the nearest N cone vectors carrying either smaller or higher *S*_p_ values. In our analysis, we have N=12. We annotated the local extremals in the 2D display of the *S_p_*(θ, φ) function to facilitate visual inspection, which is discussed below.

As shown in Figure 8(c) panel 4, we obtain a 2D display (noted as the *S*_p_-map) of the patchwise smoothness function *S_p_*(θ, φ). We perform the nearest neighbor interpolation of *S_p_*(θ, φ) on to another set of vectors defined with equally spaced θ and φ values in the spherical coordinate system representing the longitudes and latitudes respectively, and we interpolate *S_p_* values onto the new set to obtain the *S*_p_-map as shown in Figure 8(c), panel 4, where the vertical and horizontal axis are representing the longitudes θ and the latitudes φ respectively. We’d like to emphasize a fundamental prior knowledge about the *S*_p_-map: The nature of nearest neighbor interpolation yields the tiled features in the *S*_p_-map, where each tile corresponds to one cone and should represent patches with nearly identical angular areas. But in the 2D display of the *S*_p_-map, the sizes and shapes of the tiles are distorted. Such distortions are introduced by the projection from a 3D surface to a 2D map and are inevitable. However, the prior knowledge about the source of such distortions and the awareness of the actual nearly identical angular areas of the tiles in the *S*_p_-map can serve as the visual guidance to help the readers to avoid confusion, even though the tiles are shown as distorted. Future work can deploy advanced visualization methods that enable linking the patches with the 3D cell surface with interactive visualization to mitigate the distortions.

### 4.4. Polarity characterization

To characterize the polarity of the cell, we developed the polarity vector ***R***_pol_ based on the volumes of the cone-cell intersection, *V*_p_, and the set of cone vectors **C**={***r**_c_*}. When we focus on one cell, the collection of *V*_p_ values from all specific cone-cell intersection can be expressed as a set defined in the support specified by the cone vectors {*V*_p_(***r***_c_)|***r***_c_∈**C**}. Our cell polarity is derived based on the 3D interceptions of the cell with all the cones as show in Figure 9. For each cell, first we obtain an estimation of the *V*_p_ value for a cone if the cone was intercepting with a perfect sphere (indicated as the white shell in Figure 9(a)) that has the same volume of the cell. We denote this value as Ω and it is obtained by taking the average of *V*_p_(***r***_c_) across all cone-vectors.

**Fig. 9.**
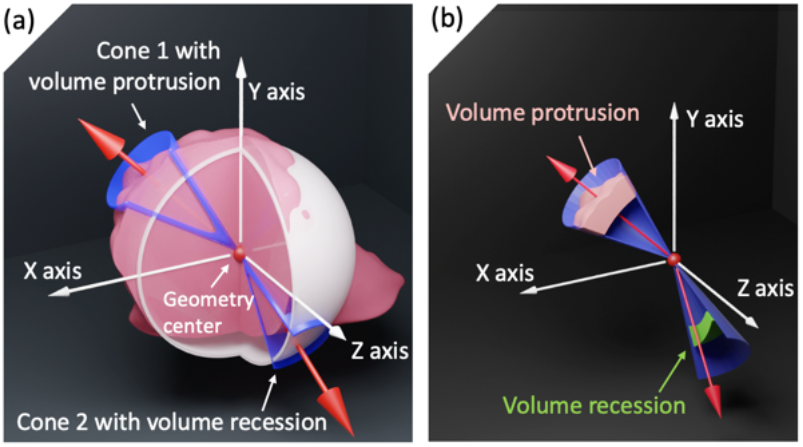
Cell polarity characterization. (a) We first define a perfect sphere that has the same volume with the cell and centered at the same geometry center shown as the while spherical shell. When the volume cropped by the cone on the cell is larger than that on the sphere as shown in Cone 1 in (a), we have volume protrusion in the angular area defined by the cone, and the value of the volume protrusion is the excess volume from the cell-cone intersect as compared to the cell-sphere intersect as annotated in (b) with the pink block. Similarly, we have volume recession indicated by the empty area shown in (a) for Cone 2 and the green block annotated in (b), and the value of the volume recession is negative. In case where both pink and green block exists for one cone, the net volume would be the final volume, which is recognized as protrusion when the value is positive, and recession otherwise.

Next, we subtract Ω from each *V*_p_(***r***_c_) to obtain the offsets of the cone-cell intersection volumes δ*V*_p_(***r***_c_) = *V*_p_(***r***_c_) - Ω. As shown in Figure 9(b), the offset represents volume protrusion when δ*V*_p_(***r***_c_)>0, and volume recession when δ*V*_p_(***r***_c_)<0. the collection of all the offsets can be expressed as a set: **{**δ*V*_p_(***r***_c_) ļ ***r***_c_ ∈ **C**}. We can define subsets for volume protrusions and volume recessions separately based on the sign of the offsets, δ*V*_p_(***r***_c_). Specifically, we obtain the protrusion set **P**={δ*V*_p_(***r***_c_) ļ ***r***_c_∈**C** and δ*V*_p_(***r***_c_)>0} and the recession set **R**={δ*V*_p_(***r***_c_) | ***r***_c_∈**C** and δ*V*_p_(***r***_c_)<0}. From set **P**, we define a protrusion center **P**_o_ by taking the geometry average of vectors defined by the cone vectors involved in **P** and weighted with the absolute value of the corresponding δ*V*_p_(***r***_c_). Similarly, we define the recession center **R**_o_ from set **R**. The polarity vector of the cell (***R***_pol_) is then defined as a vector pointing from **R**_o_ to **P**_o_. The length and direction of ***R***_pol_ characterizes the amplitude and the direction of the cell polarity respectively.

## 5. Results

We applied the analysis to all the datasets, generated a data report per imaging session to facilitate efficient inspection of the results and performed statistical comparison between the naive macrophage cells and the *Legionella* infected macrophage cells. Below we discuss the data report, the statistical comparison as implicated by the cell migration and whole-cell smoothness, as well as the statistics and the dynamics of patch-wise cell smoothness characteristics and the cell polarity.

### 5.1. Data report

Figure 10 shows a screen shot of the data report. In our data analysis pipeline, for each dataset, 46 visualization elements are produced in formats of pictures of plots, tables, images, 3D renderings, and MP4 movies, .gif files, as well as meta data information stored in HDF5 format. Browsing through the entire dataset is challenging. We acknowledge that an integrated interactive webpage tool to facilitate the data browsing and inspection would serve the purposes well, which inspires in-depth follow-up work that involves more efforts on software engineering, but in this work, we designed a series of data report as HTML pages as a lite solution to facilitate inspection of the datasets together with the analysis results. The scripts to generate such HTML pages are included in the *llsmvis* and are now openly available on GitHub [32].

**Fig. 10.**
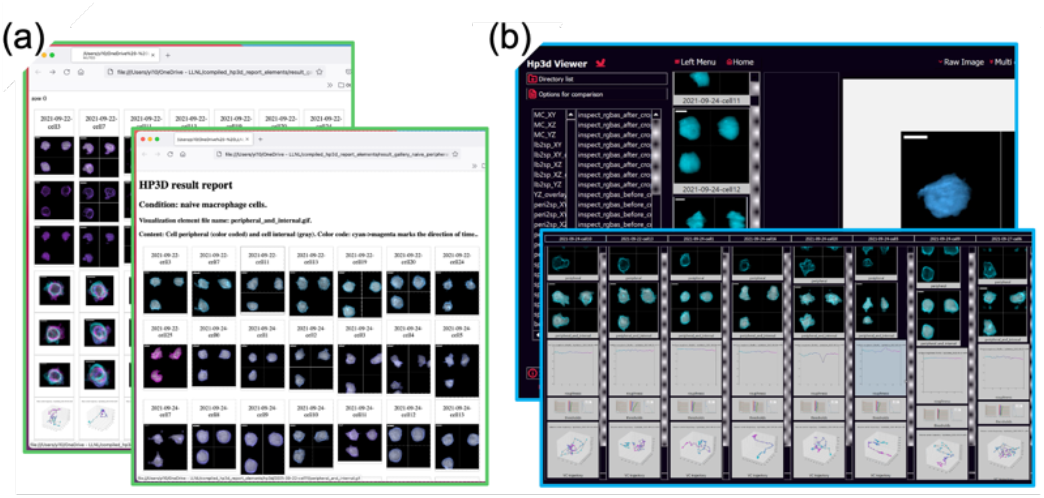
Data report screen shots. (a) Include the screen shots of html files and (b) include the screen shots of a python-based GUI, both are developed to facilitate data browsing. The HTML files can be downloaded from the data repository on figshare [31], the GUI is included in the *llsmvis* repository on GitHub [32]. Please refer to the actual *.html files and the GUI tool for better illustration and resolution of the pages. The tutorials on how to unpack the files are included in the *llsmvis* repository [32]. Scale bars used in the visualization elements are 8 μm.

### 5.2. Statistical comparison

We first compared the migration dynamics of the cells between the naïve and infected macrophage populations as shown in Figure 11(a). We use the geometric center of the cell body to represent the cell position as obtained from Figure 7(f) after the segmentation step. Each cell is imaged for 50 time points over approximately 26 minutes, so each cell exhibits a migration trajectory consisting of 50 times steps. Figure 11(a) compares the mean square displacement (MSD) of the cell trajectories as a function of the delay time (dT) to characterize cell migration behavior. Our results demonstrate that the naïve macrophage cells exhibit a slight confinement effect in the migration behavior (MSD-dT curve bends down); this agrees with the confinement imposed by the collagen matrix. Interestingly, we find a correlation of reduced confinement effect with *Legionella* infection as shown by the less-bent MSD-dT curve in Figure 11(a) for the infected macrophage cells. No directed motion is observed in the MSD-dT plot where the curve would be bent up; this agrees with our sample preparation condition where no gradient of chemoattractant is introduced to the sample.

**Fig. 11.**
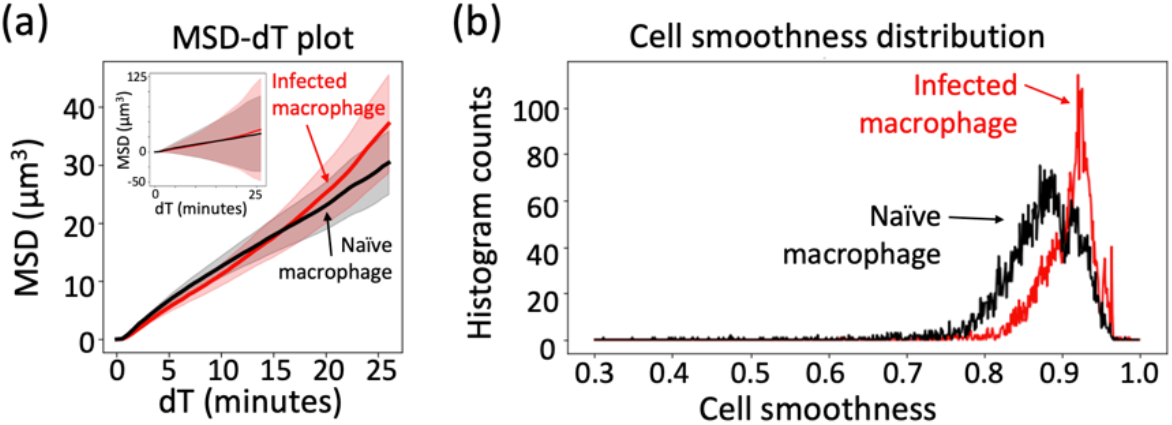
Cell migration and cell smoothness statistics. (a) shows the average mean square displacement (MSD) of the cell migration trajectories as a function of delay time (dT) (shown as red(black) solid lines), where the cell positions are defined as the geometry center of the cell at each time point. The MSD curves are the averages across all the MSD curves calculated for all the cell trajectories under the same condition. The semi-transparent pink(gray) background behind the solid red(black) lines indicates the error range. The error range shown in the main panel is defined as one Standard Error below and above the average values with SE=*d*/*n*^1/2^, where *d* is the standard deviation of the MSD at each lag time, and *n* is the number of trajectories used to obtain the average value. The error range shown in the intersect is defined as one standard deviation of the MSD values below and above the average values. Our results show that the infected macrophage cells demonstrate a free diffusion behavior in their migration patterns while the naïve macrophage cells demonstrate a slightly confined random walk pattern. The large variance shown in the intersect also demonstrates the large diversity among different cells. The infected macrophages demonstrate larger variance than the naïve macrophage cells. The migration speeds are similar between the two conditions. (b) shows the cell smoothness statistics for two conditions characterized over all the cells prepared for the condition over all time points, and the infected macrophage cells are smoother than the naïve macrophage cells.

We then calculated the whole-cell smoothness between the two different conditions (Figure 11(b)). We can see from Figure 11(b) that the *Legionella* infection is correlated with higher whole-cell smoothness.

Next, we investigated the patch-wise smoothness distribution of the cells as characterized by the number of local extrema (Figure 12). We find out that the probability histograms of the number of local extrema are similar between the naïve and the infected macrophage cells (Figure 12(a)(b)). We characterized the memory effect of such time series by calculating their autocorrelation (AC) coefficients, which is defined as the Pearson’s correlation coefficient of the time series with itself with a series of lag times (dT) as defined in Figure 13(g). For each lag time, the AC coefficients are averaged across all the 4D datasets under the same condition to yield the average AC curve. Because the Pearson’s correlation coefficient describes the similarities between the time series, the average AC curve therefore demonstrates the memory effects of the time series as indicated by its self-similarities with time offsets.

**Fig. 12.**
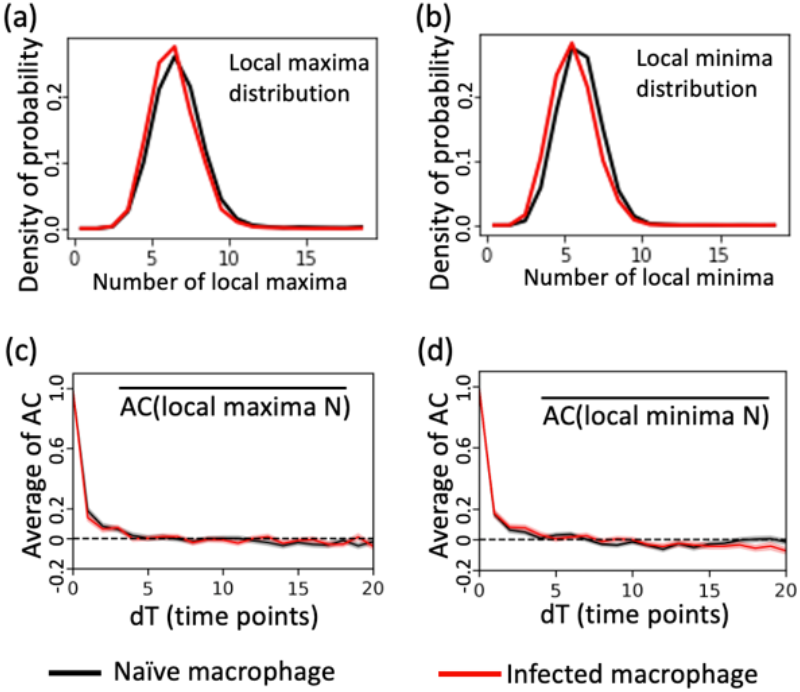
Patch-wise smoothness statistics and dynamics. Panel (a) and (b) shows the statistical distribution of the local extremals of the patch-wise smoothness profile of the naïve and infected macrophage cells, for which both exhibits similar distribution with the number of local extremals centered around 6 to 7 per cell. (c) and (d) is the averaged auto-correlation coefficients of the local extremal fluctuation profile, which demonstrate no memory effect in the patch-wise smoothness profiles for all cases. The pink(gray) background behind the solid red(black) lines indicates the error range for the average AC coefficients.

**Fig. 13.**
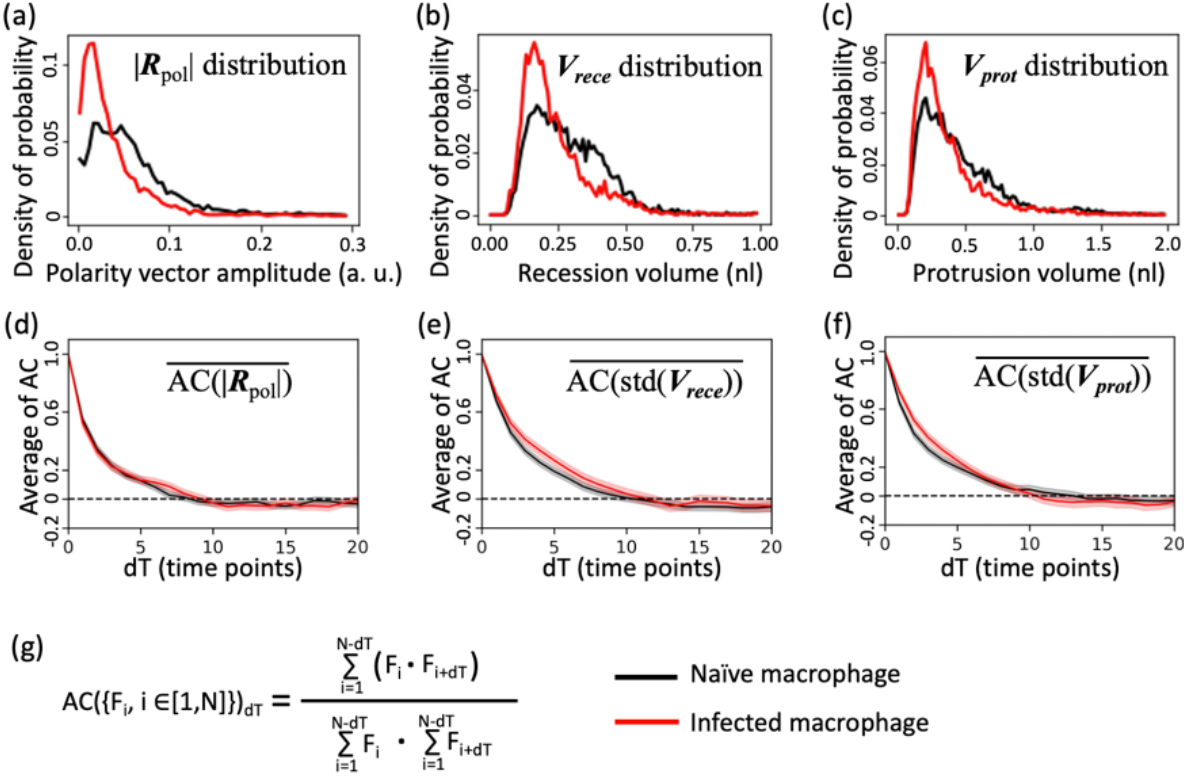
Cell polarity statistics and dynamics. (a)(b)(c) demonstrates the probability distribution of the cell polarity vector amplitudes, recession volume and protrusion volume respectively, and (d)(e)(f) demonstrates the averaged time auto-correlation (AC) coefficients of the polarity vector amplitudes, recession volume and protrusion volume respectively. The pink(gray) background behind the solid red(black) lines indicates the error range for the average AC coefficients. (g) shows the auto-correlation coefficient defined as the Pearson Correlation coefficient of the sequence with itself with an offset of dT, where dT ranges from 0 to 20. Our results indicate that the Legionella infection is correlated with reduced cell polarity, recession volume and protrusion volume, in agreement with the observed increase cell smoothness. We observed Memory effects in the dynamics of all three metrics are observed for both infected and macrophage cells as shown in (d)(e)(f). No difference in the memory effect is observed in the cell polarity between the two conditions as shown in (d), while the infection is correlate with a slightly improved memory effect in the standard deviation of the recession and protrusion volumes shown in (e) and (f) respectively. The analysis scripts and high-resolution figures are available as an example Jupyter notebook [33] on the *llsmvis* repository [32].

As shown in Figure 12(c)(d), the memory effects in both cases are weak (quick decay of the average AC curves), suggesting faster dynamics of local smoothness extrema as compared to the time resolution of our acquisition configuration under our experimental conditions. The error range is calculated as the Standard Error (SE) of the average auto-correlation coefficients: SE=*d/n*^12^, where *d* is the standard deviation of the AC coefficients at a given time lag calculated across all the measurements under the same condition, and *n* is the number of measurements used to calculate the average AC coefficients. Detailed analyses are enclosed in an example Jupyter notebook [33].

In addition, we characterized the cell polarity based on the amplitudes of the polarity vector discussed in section 4.4. for the infected and naive macrophage cells. As shown in Figure 13, *Legionella* infection is correlated with reduced cell polarity (Figure 13(a)). We also calculated the averaged AC coefficients of the cell polarity vector amplitudes (Figure 13(d)), and find out that the polarity fluctuation both exhibit memory effect, suggesting that the time resolution of our acquisition routine is sufficient to capture the polarity dynamics. The memory effects for the cell polarity vector amplitudes are similar between the infected and native macrophages (Figure 13(d)), suggesting negligible differences in the dynamics of the cell polarity centers under the conditions explored in our study. We also investigated the standard deviations of the cell volume recessions (Figure 13(b)) and cell volume protrusions (Figure 13(c)) and found a correlation of reduced standard deviations (less dispersive distribution) with *Legionella* infection, suggesting less dispersive membrane configuration in infected macrophage cells. The memory effect of such standard deviations (Figure 13(e)(f)) are also observed, indicating that the *Legionella* infection is correlated with slightly increased memory effect, suggesting slower dynamics in the membrane protrusions.

All the statistical characterizations shown above are performed from 131 cells for naïve macrophage cells, and 101 cells for infected macrophage cells. We removed the invalided numbers (NaN) from all the time series when calculating the average AC curves and the corresponding error ranges, the percentage of removed data are both below 1% as shown in the Jupyter notebook for analyses [33]).

Under the conditions explored in this study, we find that the *Legionella* infection is correlated with increased cell smoothness and with migration pattern closer to random walk, while the naïve macrophage cells exhibit confined diffusion behavior that demonstrates the confinement effect imposed by the collagen matrix. The infection is also correlated with reduced amplitude of cell polarity, in agreement with the prior studies where the *Legionella* infection can cause cleavage of actin stress fibers and cause cell-rounding phenotype in HEK293T cells [27]. We also find that the dynamics of the cell polarity amplitude change is similar between naïve and infected macrophage cells, while the memory effect for the standard deviations of the volume protrusion and volume recessions are increased in infected macrophage cells, suggesting a correlation between *Legionella* infection and decreased membrane configuration dispersiveness dynamics. We acknowledge that the differences are small, motivating more rigorous biological studies in the follow up studies. The *Legionella* infection performed in our study does not influence the number of local extremals characterized by the patch-wise cell surface smoothness statistics and dynamics, where both demonstrate the number of extremals centered in the range of 6 to 7 without memory effect.

## 6. Discussion and future direction

In this work, we built and deployed a lattice light sheet microscope for the study of *Legionella* infection through novel multidisciplinary approaches optimized and systematically integrated over (a) instrument configuration, maintenance, and operations, (b) sample preparation and data acquisition design, and (c) the analysis method and pipeline. To the best of our knowledge, our work is the first deployment of LLSM to study bacterial infection. We have demonstrated that the workflow can deliver adequate throughput with the effort level attainable by a small research team. We derived quantitative figures of merit to measure the cell smoothness, cell migration and cell polarity characteristics to characterizes our datasets; we also characterized the statistics and dynamics for both the infected and naïve macrophage cells each over more than 100 cells. Our results provide new insights into the *Legionella* infection study in a novel observation domain enabled by LLSM that is previously inaccessible. Additionally, our work facilitates further focused-technical development and biological explorations by removing the interdependencies between the aspects of instrumentation, sample, and data, providing directions for further advancements in each direction independently to further utilize LLSM for the subject of study. For example, in the instrumentation aspect, further automation of the data acquisition routine is plausible and can be adopted by independent commercial LLSM providers or the opensource software developer groups such as μManager [28] and pycro-manager [29]. In the sample preparation aspect, focused molecular biology experiments can be designed and explored without extensive involvement of LLSM imaging to push the research further into specific molecular mechanisms studies in integration with more traditional molecular biology approaches. In the data analysis and visualization aspect, further advancements in the surface morphometrics characterization algorithms, data management and data visualization platforms can be performed, and supported by (a) the datasets collected in this study, and (b) our analysis scripts which serve as an adaptor to access LLSM output datasets for fast prototyping. The analysis can be adopted by open-source microscopy data analysis tool set such as *napari* [30]. We expect our work to promote the adoption of LLSM with a workflow suitable for both the academic research teams and the commercial providers and users of LLSM.

## Supporting information

visualization 1

visualization 2

visualization 3

visualization 4

visualization 5

visualization 6

## 7. Author contributions

X. Yi designed and executed the study with inputs from the whole team. T. A. Laurence and X. Yi designed and built the modified LLSM. X. Yi developed the LLSM alignment, maintenance, and imaging workflow. X. Yi calculated and implemented the SLM patterns used in this study. X. Yi designed and performed the biological experiments with guidance from K. W. Overton and inputs from J. K. Lo, who performed on-demand *Legionella* preparations, and T. H. Lee, who performed preparatory *Legionella* culture work. X. Yi designed the data analysis methods and implemented the analysis pipeline with guidance from P. T. Bremer and inputs from H. Miao, who implemented cell surface extraction and cone-cropping operations. Y. Zhang implemented the Python based graphical user interface to inspect the visualization elements generated in this study. K. W. Overton and X. Yi made the mCherry expressing RAW264.7 cell line. M. M. Elsheikh made the mCherry expressing *Legionella*. C. Jiang implemented the 3D cell animation generation pipeline, and advised on 3D graphics rendering. B. W. Segelke advised on microbiology. K. W. Overton advised on mammalian cell culture and engineering. P. T. Bremer advised on data analysis and high-performance computing. T. A. Laurence advised on instrumentation. B. W. Segelke, K. W. Overton, P. T. Bremer and T. A. Laurence supervised the study.

## Acknowledgement

We thank Dr. Srigokul Upadhyayula and Dr. Eric Betzig for the discussions on the imaging workflow and for the support of test imaging samples for LLSM. We thank Dr. Dina R. Weilhammer for supplying the RAW264.7 cell line, and we thank Dr. Grant J. Jensen for sharing the *Legionella* bacteria [5]. We also thank Dr. Yuxi Liu and Ms. Jue Wang for discussions on *Legionella* culture and infection experiments and the support on *Legionella* bacteria culture materials. We thank Dr. Rikki Garner for discussions on collagen preparation. We thank Dr. Loic Royer, Dr. Bin Yang, and Dr. Bo Huang for general guidance with microscopy and data analysis. We thank Dr. Howard Shuman for advice and discussions on *Legionella* pathogenesis. We also thank Dr. Dante Paul Ricci for discussions on microbiology, and Dr. Dan M. Park with electroporation experiments. Work at LLNL was performed under the auspices of the US Department of Energy under contract DE-AC52-07NA27344. Release number LLNL-JRNL-831386.

## Data availability

The datasets are now openly available online as the dataset repository *“Datasets for the manuscript titled “A Tailored Approach to Study Legionella Infection Using Lattice Light Sheet Microscope (LLSM)””* on figshare [31].

## Appendix

**Fig. A1.**
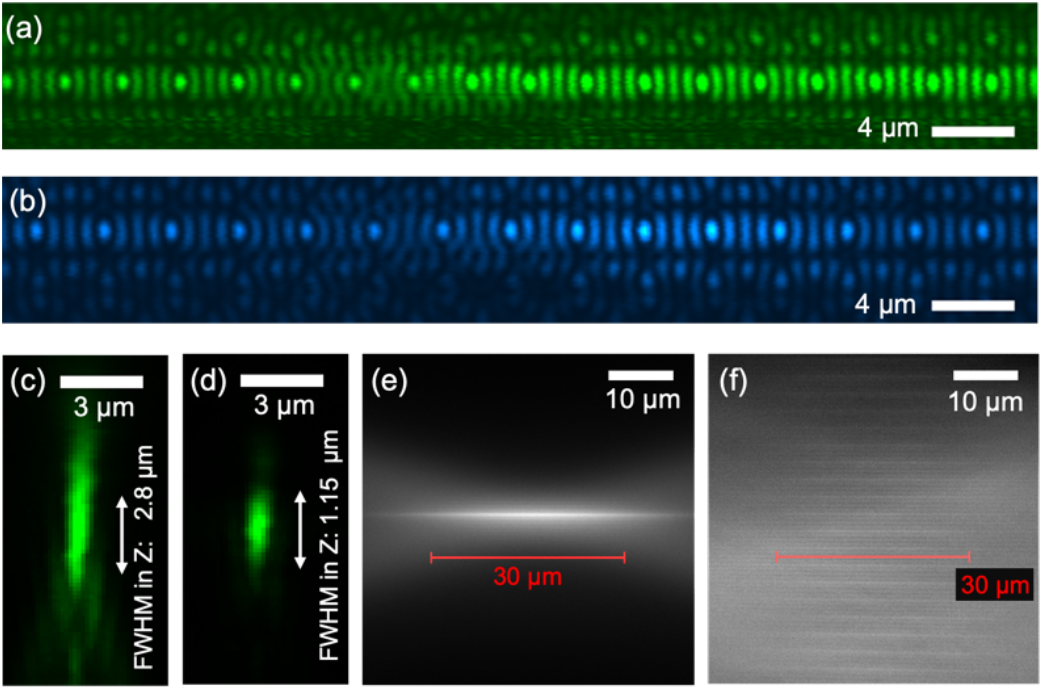
PSF and light sheet characterization. (a) and (b) demonstrate the experimentally measured light lattice cross-section intensity at the sample for 561 nm and 488 nm excitation respectively. The measurement is performed by scanning a bead across the XZ cross-section and take the integral of intensities to represent the excitation field intensity at each position. (c) and (d) demonstrate the XZ PSF measured with a green bead excited by the 488 nm light sheet for the detection PSF (measured by scanning the detection objective without scanning the light sheet) and overall PSF (measured by co-scanning the detection objective and the light sheet along the axial direction of the detection path) respectively. (e) and (f) demonstrate the experimentally measured 488 nm excitation profile of a single Bessel beam and a light lattice respectively. The measurements were performed by imaging fluorescein solution with the corresponding excitation light field. The solution is made with 100 μl saturated fluorescein solution mixed in 7 ml of distilled water.

